# Cortical recruitment determines learning dynamics and strategy

**DOI:** 10.1101/274936

**Authors:** Sebastian Ceballo, Jacques Bourg, Alexandre Kempf, Zuzanna Piwkowska, Aurélie Daret, Thomas Deneux, Simon Rumpel, Brice Bathellier

**Affiliations:** Unité de Neuroscience, Information et Complexité (UNIC), FRE 3693, Université Paris-Saclay and Centre National de la Recherche Scientifique, Gif-sur-Yvette, F-91198, France; Institute of Physiology, Focus Program Translational Neuroscience, University Medical Center, Johannes Gutenberg University, Mainz, Germany

## Abstract

Salience is a broad and widely used concept in neuroscience whose neuronal correlates, however, remain elusive. In behavioral conditioning, salience is used to explain various effects, such as stimulus overshadowing, and refers to how fast and strongly a stimulus can be associated with a conditioned event. Here, we show that sounds of diverse quality, but equal intensity and perceptual detectability, can recruit different levels of population activity in mouse auditory cortex. When using these sounds as cues in a Go/NoGo discrimination task, the degree of cortical recruitment matches the salience parameter of a reinforcement learning model used to analyze learning speed. We test an essential prediction of this model by training mice to discriminate light-sculpted optogenetic activity patterns in auditory cortex, and verify that cortical recruitment causally determines association or overshadowing of the stimulus components. This demonstrates that cortical recruitment underlies major aspects of stimulus salience during reinforcement learning.

## Introduction

Sensory stimuli can vary substantially in their efficacy to serve as a conditioned stimulus during behavioral conditioning. In the context of classical conditioning, a well-known example is the so called “overshadowing” effect. When animals are trained to associate two simultaneously presented stimuli (historically a tone and a flash) to a specific unconditioned stimulus (e.g. foot-shock), it is often observed that, after training, the animal is conditioned more strongly to one stimulus than to the other^1,2^. In their seminal theoretical work originally developed for classical conditioning, but later extended to operant conditioning (e.g. in reinforcement learning), Rescorla and Wagner^3^ introduced the notion of salience to explain the overshadowing phenomenon. In their model, salience is a parameter affecting the speed at which a given stimulus is associated with the unconditioned stimulus. Thus, when behavior reaches maximal performance and learning stops, the more salient of the two stimulus representations has been associated more strongly with the unconditioned stimulus, leading to overshadowing. While this theory captures a number of phenomena and is the basis for important frameworks such as reinforcement learning^4,5^, the neural underpinnings of the salience parameter remain elusive.

Salience in this context is usually seen as the global amount of neural activity representing the stimulus, like in “pop out” models of attentional salience^6–8^. This intuitively follows from the idea that if more spikes are involved in representing a stimulus, they can produce more synaptic weight changes, as expected from the firing rate sensitivity of typical learning rules^9–13^, and thus modulate more rapidly the relevant connections. However widespread, this idea lacks direct causal experimental verification in a learning task. Moreover, other theories propose that salience could also be encoded in other parameters such as neuronal synchrony levels^14–17^, which could influence learning via the temporal properties of biological learning rules^18–22^. Thus, the neuronal correlate of stimulus salience is a key question with broad implications for learning theories.

Using auditory discrimination tasks of sounds with different global cortical response strengths, we show that cortical recruitment impacts learning dynamics^23,24^ in a manner similar to the salience parameter of a reinforcement learning model. To explore this result in more precise experimental settings, we trained mice to discriminate optogenetically-driven response patterns that elicit different levels of cortical activity. Using this paradigm, we directly demonstrate that cortical recruitment determines which part of a compound stimulus drives a learned association while “overshadowing” other parts of the stimulus. This validates a generic prediction of reinforcement learning models and causally establishes the role of cortical recruitment as a neuronal correlate of stimulus salience.

## Results

### Sounds with identical physical levels and perceptual detectability can recruit different levels of cortical activity

It is often assumed that i) the physical intensity of a stimulus determines ii) the amount of recruited neuronal activity in the brain and iii) its perceptual detectability, which represents the salience of a particular stimulus^25^. We thus wondered how tightly these three properties are related for sounds, taking into consideration the nonlinearities reported for the encoding of sound features^26,27^. We chose three short, complex sounds (70ms duration) containing a large range of frequencies and temporal modulations (**Fig. 1a**). The sounds were normalized to have equal mean pressure levels (standard deviation of the waveform). We first tested whether this set of normalized sounds are also equally detectable by mice in a behavioral task. We initially trained mice to lick on a water port after presentation of each of the three sounds (all mice experienced the three sounds in the same task) to obtain a reward (**Fig. 1b**). We then measured behavioral response probability to decreasing intensity levels of the sounds. We observed that, for all sounds, response probability steadily decreased down to chance level as measured in the absence of sound (**Fig. 1c**). Yet, no significant difference in response probability curves was observed across the three sounds (Friedman test – non-parametric anova, p = 0.43, n=6), indicating that, in this context, detectability was mainly related to sound intensity and not quality.

**Figure 1:**
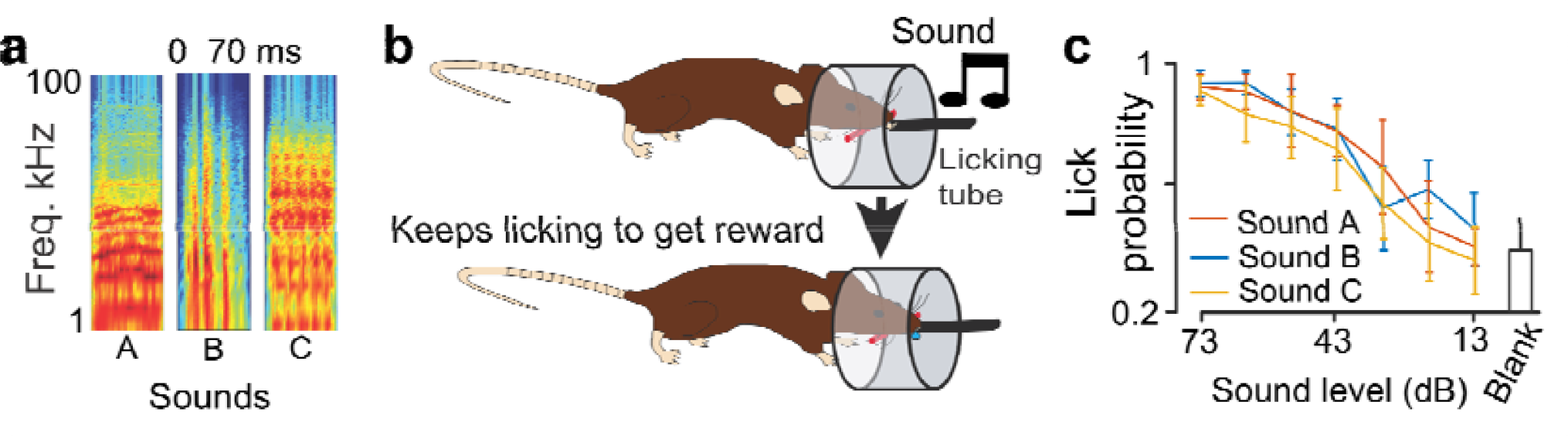
Spectro-temporal differences in complex sounds do not affect near-threshold detectability. ***a**. Spectrograms of three 70 ms long complex sounds. **b**. Schematics describing the auditory detection task. **c**. Mean response probability for 6 mice trained to detect sounds A, B and C at 73 dB to get a reward and probed with lower sound intensities. While the effect of intensity was significant, there was no effect of sound identity (Friedman test, p_intensity_ = 2.3×10^−9^, p_sound_ = 0.43, n=6 mice)*.

We next wondered whether recruitment of neural activity in the auditory cortex in response to these three sounds was also independent of sound quality and determined by the sound level. To do so, we used two-photon calcium imaging in awake, passively listening mice, a technique that offers access to large samples of neuronal activity. We imaged 6 mice that were injected with AAV1-GCAMP6s virus in auditory cortex (**Fig. 2a**). Recordings were followed by an automated image registration and segmentation algorithm (**Fig. 2b**)^28^ that allowed the isolation of 15,511 neurons across 27 imaging sites, from which large fluorescence signals could be observed (**Fig. 2c**). The fields-of-view were either 0.5×0.5 or 1×1 mm (**Fig. 2a, b**), allowing a rapid tiling of the full extent of primary and secondary auditory cortex (**Supplementary Fig. 1**). Cortical depths were randomly chosen ranging between 100 and 300 µm corresponding to layer 2/3. The mouse auditory cortex (primary + secondary) contains approximately 200,000 neurons in one hemisphere^29^ and thus about 50,000 neurons in layer 2/3, so we expect our sample of ~15,000 neurons to be representative for supragranular auditory cortex. Comparing the amplitude of the mean-deconvolved calcium signals recorded across the entire duration of the response (see example population response profiles in **Fig. 2d**), we observed that at 73 dB intensity, sound A elicited at least two-fold less cortical activity than sounds B and C (0.05 against 0.10 and 0.12 % ΔF/F.s^−1^, **Fig. 2e**). This was consistently observed across mice (**Supplementary Fig. 1**). Furthermore, sound A triggered a significant response (Wilcoxon sign test across 20 sound repetitions) in only about 35% of all neurons, while sounds B and C significantly excited 41% and 45% of all neurons (**Fig. 2e**). Furthermore, this difference in population responsiveness was consistent with previous, independent measurements performed under anesthesia (**Supplementary Fig. 1**)^30^. It is noteworthy that all three sounds elicited distinct response patterns as evaluated by correlation-based population similarity measures. Thus, sound identity could be decoded with high accuracy, on a single-trial basis, using linear classifiers (**Fig. 2f**). This shows that sound discriminability was not affected by cortical recruitment.

**Figure 2:**
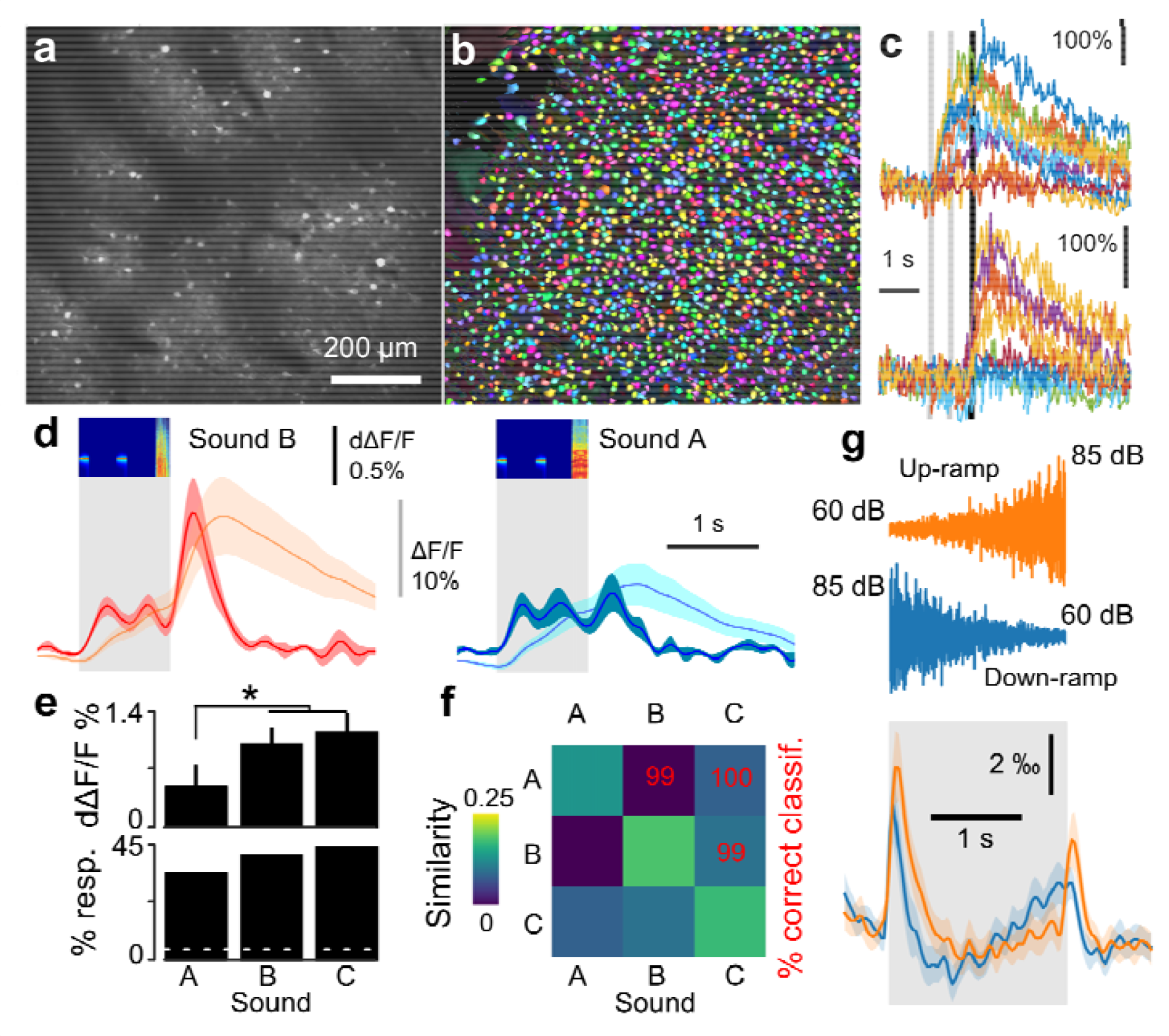
Spectro-temporal differences impact on cortical recruitment. ***a**. Example field of view illustrating GCAMP6s labeling of L2/3 auditory cortex neurons. **b**. Result of automated cell segmentation run on the data acquired in the example shown in **a**. **c**. Example single trial responses to sounds (different colors) for two neurons (top and bottom). **d**. Population responses (n = 26 sessions, 15511 neurons in 6 mice) to Sound B (red) and A (blue). Both normalized fluorescence (light colors) and deconvolved (dark colors) calcium signals are shown. **e**. Mean deconvolved signal and fraction of significantly responding neurons (Wilcoxon signed test, p<0.05) to sounds A, B and C. Mean calcium responses to sound A (0.05 ± 0.03% ΔF/F.s^−1^) were significantly smaller than to B (0.10 ± 0.02% ΔF/F.s-^1^) and C (0.12 ± 0.02% ΔF/F.s^−1^; sign test, p = 0.0094 and p = 0.029, n = 26 sessions, 15511 neurons in 6 mice). The fractions of responding neurons (35, 41 and 45 %) were all significantly different from each other (χ^2^ test, p = 10^−12^, 10^−27^ and 3×10^−72^, n = 15511). **f**. Population response reliability (diagonal) and similarity (off-diagonal) matrix for sounds A, B and C. The pair-wise discriminability value, computed with a linear classifier is indicated in red. **g**. Mean deconvolved calcium signals for 6757 auditory cortex neurons in 12 awake mice during 29 calcium imaging sessions for 2 s long white noise sounds modulated in intensity between 60dB to 85dB upwards and downwards*.

Another discrepancy between cortical recruitment and the physical intensity of a stimulus can be observed using sounds with different temporal intensity profiles. Up-ramping sounds elicit larger cortical responses in mice^26^ and other animals^31,32^ than their time-symmetric down– ramps, despite the equality of their cumulative physical energies. This effect correlates with asymmetries in subjectively perceived loudness in humans^33,34^. We confirmed this result (**Fig. 2g**) for 2s white noise sounds ramping between 60 and 85dB (mean deconvolved ΔF/F measured from 0 to 2.5 s after sound onset, up: 0.166 ± 0.58 ‰ vs down: 0.157 ± 0.57 ‰, Wilcoxon rank-signed test, p = 0.034), with a clear effect even at the onset despite the lower start intensity level in up-ramps (mean between 0 and 0.5s for up: 0.80 ± 0.08 ‰ vs down: 0.62 ± 0.11 ‰, Wilcoxon rank-signed test p=3.75×10^−4^, n = 29 sites, in 12 mice).

In summary, although intensity is obviously crucial for detectability, when sounds are played at an intensity above the detection threshold, the specific amount of recruited cortical activity strongly depends on factors other than intensity and can vary across different sounds. Intrigued by this observation, we asked whether cortical recruitment could be related to stimulus salience in a learning task.

### Cortical recruitment correlates with learning speed

Classically, relative salience measures are performed using an overshadowing paradigm in which two stimuli (e.g. a flash and a sound) are conditioned together, as a compound stimulus, to an unconditioned stimulus (e.g. foot-shock). Then, salience is derived from the level of the conditional response elicited by each stimulus component individually. While this approach is valid when the compound is made of stimuli from two different sensory modalities, two simultaneous sounds are likely to fuse perceptually, precluding measurement of their individual saliences with the classical overshadowing design^35^. Thus, we measured stimulus salience through learning speed, using an auditory-cued Go/No-Go task. To do so, water-deprived mice were first trained to visit a lick-port and to receive a water reward if they licked after being presented with an S+ sound. This pre-training phase mainly aimed at raising motivation in all mice and was not used to measure learning speed. When mice collected rewards in at least 80% of their port visits, the Go/NoGo task was started by introducing a non-rewarded S- sound in half of the trials (**Fig. 3a**). After a large number of trials, mice succeeded to both sustain licking to the S+ and withdraw from licking for the S- (**Fig. 3b**), thereby demonstrating their ability to discriminate the two sounds. Importantly, as typically observed in such tasks^24^, the S+ sound was rapidly associated with the lick response and the rate limiting factor in the acquisition of the task was to associate the suppression of licking with the S- sound (**Fig. 3b**). Hence, learning speed depends more on the salience of the S-than of the S+ sound in this task. The relative saliences of two stimuli X and Y can thus be measured by comparing the learning speed of the X versus Y Go/NoGo discrimination when X is the S- against the speed observed when Y is the S-. If X recruits less activity than Y, we expect learning to be slower when X is the S-. We therefore trained six cohorts of mice to compare the salience of sounds pairs A-B, A-C and B-C, and we used data from an earlier study to compare salience of up and down-ramping sounds^26^. Plotting the population learning curves for the sound pairs with maximum cortical recruitment differences (A-B & A-C,), we qualitatively observed that the average learning speed was faster when cortical recruitment for the S- sound was larger than for the S+ sound (**Fig. 3c**), suggesting a positive correlation between salience observed via learning speed and cortical recruitment.

**Figure 3:**
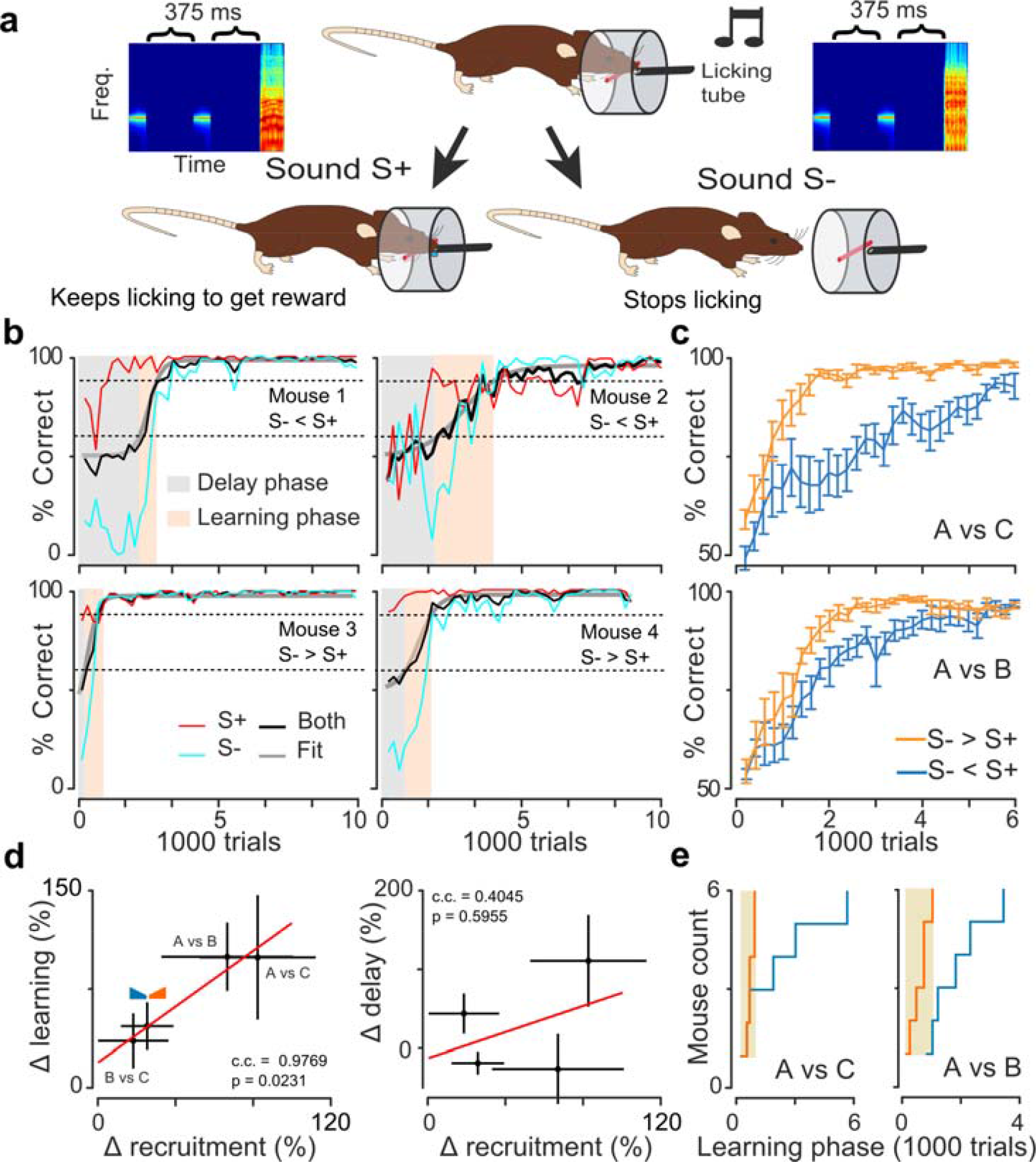
Learning phase duration correlates with cortical recruitment differences. ***a**. Schematics describing the auditory Go/NoGo discrimination task. **b**. Individual learning curves for 6 mice discriminating sounds A and C. Performance for S+ (red), S- (light blue) and both (black) sounds are displayed. Mice from the top row have sound A as the S- stimulus while mice from the bottom row have sound C as the S- stimulus. Typical learning curves display a delay and learning phase as shown in light gray and orange colors. **c**. Mean learning curves for different groups of mice (n = 6 for each curve) discriminating between sounds A and C (top) or A and B (bottom). Slower learning is observed when the S- sound recruits less cortical activity than the S+ sound (blue) as compared to when sound valence is swapped (orange). **d**. Mean +/- standard error of the difference of learning (left) and delay (right) phase durations plotted against the difference in neural recruitment by the stimuli. A significant correlation coefficient of 0.9769 (p=0.0231, n=4) is observed for the learning phase, while the weaker correlation of 0.4045 for the delay phase is not significant (p=0.5955, n=4). **e**. Cumulative distributions of learning phase durations for sound pairs A-C, A-B*.

However, looking at learning curves from individual mice, we noticed that the qualitative difference observed at the group level hides a more complex effect. As often observed in animal training^23^ and as we previously reported for the particular task used in this study^24^, most individual learning curves had a sigmoidal rather than exponential time course. Specifically, the curves displayed a delay phase with no increase in performance followed by a learning phase with an often very steep performance increase. Also, the duration of each phase was highly variable across animals as exemplified in **Fig. 3b**. We wondered whether cortical recruitment was affecting one particular phase or both. We performed a sigmoidal fit (**Fig. 3b**) on each individual curve from which the delay phase duration was computed as the number of trials necessary to reach 20% of maximal performance level, and the learning phase duration as the number of trials necessary to go from 20% to 80% maximal performance. We then plotted the difference of cortical recruitment for the two discriminated sounds against the observed difference in learning or delay phase durations when one or the other sound is S- (**Fig. 3d**). We observed a significant positive correlation between cortical recruitment and learning phase duration but no correlation for the delay phase (**Fig. 3d**). These results were corroborated by an analysis of variance showing that the sign of the difference between recruitment for S+ and S- has a significant effect on learning phase duration (**Supplementary Fig. 2**). In addition, we noticed that cortical recruitment had an effect not only on the mean duration of the learning phase at the group level but also on the inter-individual variability. When the S- sound recruited more activity than the S+ sound, learning phase duration was more homogenous than for the opposite sound assignment, especially for the two sound pairs with a large difference of cortical recruitment (**Fig. 3e, Supplementary Fig. 2**).

### A reinforcement learning model captures the effects of cortical recruitment on learning dynamics

In order to better understand the observed experimental correlation between cortical recruitment and learning dynamics, we employed a recently developed model of the discrimination task. This model is based on the Rescorla-Wagner reinforcement learning framework but extending it to a simple but more biologically interpretable model (**Fig. 4a**)^24^. In short, the model postulates that associative learning occurs by adjusting the synaptic weights between “sensory” and “decision” neural populations described by population firing rate variables. At the input, two populations are specific for the S+ and S- sounds respectively, which we denote as Ŝ+ and Ŝ-, and one population, Ĉ, which represents information common to S+ and S- trials (e.g. overlap between the S+ and S- representations or activity independent of sound, for example, related to visiting the lick port). Population Ĉ is an essential element of the model to reproduce high initial ‘hit’-rates combined with delayed discrimination learning^24^. The “decision” population has two ensembles: one promoting and one inhibiting licking (**Fig. 4a**). Adjustment of synaptic weights happens through a Hebbian learning rule modulated by Rescorla and Wagner’s (1972) δ-rule which gates weight updates by the reward expectation error. However, the employed δ-rule is asymmetric, meaning that the learning rate is larger by a factor *v* when an unexpected reward occurs, as compared to when an expected reward does not occur. This asymmetry is crucial for capturing the asymmetry of the learning process (i.e. the fast adjustment of the ‘hit’-rate, and slower adjustment of the ‘correct rejection’-rate; **Fig. 3b**). In addition, synaptic updates are multiplicative, meaning that weight updates are proportional to the current weight value^36–39^. The key feature of multiplicative learning is that learning speed depends on the current strength of the synapses. Thus, the same model can have very slow learning (low weights) as in the delay phase and faster learning (high weights) as in the learning phase. Furthermore, this feature makes learning dynamics highly sensitive to the initial synaptic weights, which become important parameters that can even account for most of the inter-individual variability^24^. Most importantly here, the model has an explicit salience parameter, that corresponds to the activity of the Ŝ+ and Ŝ-populations, which makes it well-suited to interpret our experiments.

**Figure 4:**
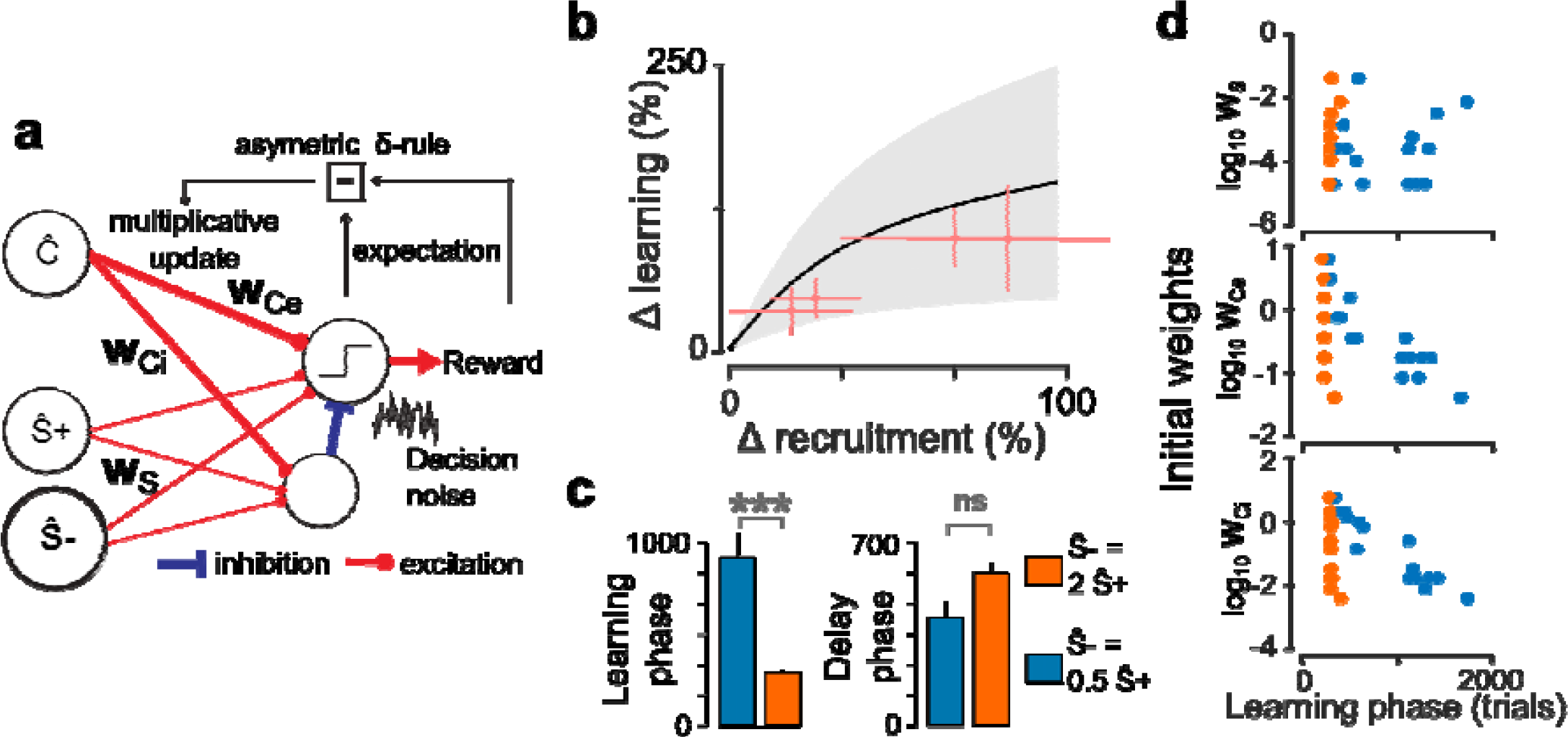
A multiplicative reinforcement learning model explains salience-related variations in learning speed. ***a**. Schematics describing the auditory Go/NoGo discrimination model. **b**. Mean +/- standard deviation of the difference of learning phase duration against difference in neural recruitment by the stimuli for the model initialized with 15 initial conditions obtained by fitting learning curves from a previous study^24^. The experimental observations of **Fig. 3d** are superimposed in red. **c**. Mean +/- standard errors for the learning and delay phases obtained with the model for a two-fold difference in cortical recruitment between the two stimuli. A longer learning phase (Kolmogorov-Smirnov test, p=8×10^−7^, n=15 initial conditions per group) but not delay phase (Kolmogorov-Smirnov test, p=0.13, n=15 initial conditions per group) is observed when S- recruits less activity (blue) as compared to when S- recruits more activity than S+ (orange). **d**. Learning phase duration plotted against the value of the modeled initial synaptic weights. Same simulations and color code as in **c**. Significant correlations were observed only when S- recruits less activity (blue), and for **w_Ce_** (ρ = −0.60, p = 0.016, n=15) and **w_Ci_** (ρ = −0.68, p = 0.005, n=15)*.

We therefore wondered whether this model reproduces the observed effects of cortical recruitment on mean learning speed and its variability. The dynamics of the model depend on the choice of its three core parameters (noise level, learning rate and asymmetry, see Methods) and of its initial synaptic weights, which could be fitted from the data, however with potential overfitting issues. In order to rule out these issues, we looked at the qualitative behavior of the model using a set of parameters obtained in a previous group of experiments^24^ by fitting the individual learning curves from 15 mice trained in a task identical to the one used in this study. This parameter set included 15 different values of the initial weights, and core parameters were identical for all mice, which we showed is sufficient to account for inter-individual variability^24^. Based on these parameters, and systematically varying the salience values in simulations, we observed that salience differences were positively correlated to learning phase duration and variability in the model, similar to the experimental results (**Fig. 4b**). Also, as illustrated when salience is doubled for one of the two stimuli, the model reproduced two other experimental observations. First, the delay phase was not significantly influenced by salience (**Fig. 4c**). Second, the variability of the learning phase duration was much stronger when S- recruited less activity than S+ (**Fig. 4c,d**). Thus, without any tuning, the model qualitatively reproduced the complexity of the experimental dynamics. This model thus provides a testing ground for important factors influencing the complex effects of stimulus salience on behavior in a precise theoretical framework.

### Learning speed effects related to neuronal recruitment crucially depend on initial synaptic strengths in the model

To understand the origin of salience-based effects in our simulations, we plotted the learning phase duration against the values of the three initial synaptic weight parameters (**w_Ce_** and **w_Ci_** for the Ĉ population, and a single initial value **w_S_** for the four weights of the Ŝ+ and Ŝ-population, as in **Fig. 4a**). First, we observed that the initial weight between sound-specific neural populations and the decision populations (**w_S_**) had no correlation with learning speed duration. This was expected as **w_S_** mostly impacts the delay phase and not the learning phase. Indeed, because of the multiplicative rule, small initial synaptic weights lead to slow initial learning, creating a period in which the performance does not improve above noise (delay phase). When sufficient learning has occurred (end of delay phase), the sound-specific synaptic weights become large enough to increase performance at a high learning speed. At this stage, the initial weight value is virtually “forgotten” and does not influence learning speed anymore. Our earlier fitting results^24^ showed that **w_S_** is the main determinant of the delay phase duration and is highly variable across mice. In our simulations, inter-individual **w_S_** variations induced large variations of delay phase duration masking the smaller impact of salience on delay phase. Thus, our model suggests that the independence between cortical recruitment and delay phase duration in our experiments is due to inter-individual variability in initial connectivity.

A second observation was that the learning phase duration is large when the initial weights **w_Ce_** and **w_Ci_** (non-specific population Ĉ) are small, but only when S- recruits less activity than S+ (**Fig. 4d**). In contrast, for large weights, the salience differences between S+ and S- have no effect on learning phase (**Fig. 4d**). Thus, our model suggests that the Go/NoGo task does not systematically reveal the influence of salience on learning speed, which the model predicts to appear only in some animals, depending on their initial synaptic state. This phenomenon could explain the large variability observed when cortical recruitment for S- is smaller than recruitment for S+ (**Fig. 3e**).

To better understand why the initial conditions of **w_Ce_** and **w_Ci_** gate the influence of stimulus salience on learning speed, we plotted the time-course of both excitatory and inhibitory connection weights for four combinations of salience and initial weights values (**Fig. 5a-d**). The performance plots illustrate that learning phase is strongly prolonged by lower S- salience (as compared to higher S- salience) only when **w_Ce_** and **w_Ci_** are small (**Fig. 5d**). The synaptic plots show that this large prolongation of the delay phase is due to the low initial weights from both the Ŝ- and Ĉ to the NoGo decision population. These low weights are maintained during the delay phase and because of the multiplicative rule strongly slow down learning. The performance of correct rejection responses to S- takes longer and flattens the overall learning curve. In contrast, when synaptic weights from the Ĉ population to the NoGo population are initially high, more rapid S- rejection learning can be obtained solely based on the Ĉ common population (**Fig. 5c**).

**Figure 5:**
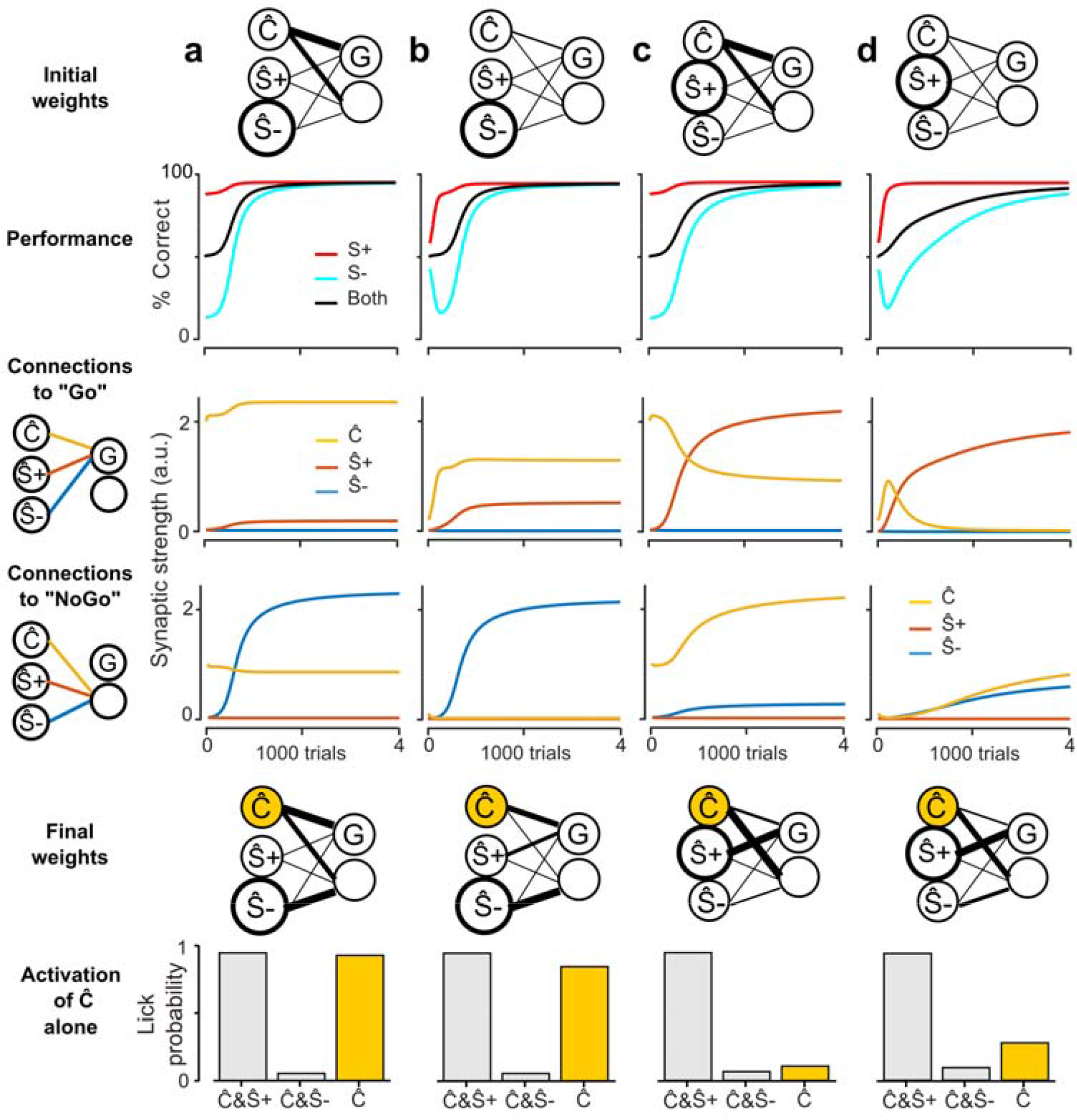
Selection of learning strategy depends on neuronal recruitment in the model. ***a**. (top) Sketch of the initial synaptic weights and simulated model performance for S+ (red), S- (light blue) and both stimuli (black). (middle) Values of the connections to the excitatory (G) and inhibitory decision populations as indicated by the schematics on the left-hand-side. Yellow: connections from the “common” Ĉ population. Red: connections from the Ŝ+ population. Blue: Connections from the Ŝ- population. (bottom) Sketch of the connectivity pattern after and response probability after learning for the S+ (co-activation of Ŝ+ & Ĉ) and S- (co-activation of Ŝ- and Ĉ) reinforced stimuli as well as for the common stimulus component alone (Ĉ, yellow). Simulation parameters: x⃗ = [1; 0.66; 1.33], α = 0.01, σ = 0.6195, v = 6, **w**_Ci_ = 1, **w**_Ce_ = 2, and **w**_S_ = 0.01. **b**. Same as **a**, but with **w**_Ci_ = 0.1, **w**_Ce_ = 0.2. **c**. Same as **a**, but with x⃗ = [1; 1.33; 0.66]. **d**. Same as **c**, but with **w**_Ci_ = 0.1, **w**_Ce_ = 0.2*.

When S- has higher salience, learning is always fast (**Fig. 5a-b**) because, in all cases, the rate limiting process remains the abolition of licking to the NoGo stimulus (due to learning rule asymmetry, see **Supplementary figure 3**). This process is boosted by strong Ŝ- salience. Also, in these conditions, acquisition of the NoGo stimulus is independent of **w_Ce_** and **w_Ci_**, because the Ĉ population drives the Go response. Thus, the refined analysis of the model suggests that a complex modulation of learning phase duration by salience is facilitated by a non-trivial assignment of the three “sensory” populations to either Go or NoGo responses, based on the salience distribution. Specifically, when the Ŝ- population is more salient, it is assigned to NoGo, while the Ĉ population drives the Go response. In contrast, when the Ŝ+ population is more salient, it is assigned to Go, and in this case, the Ĉ population drives the NoGo response.

This provides a very general and testable prediction of reinforcement learning models using population activity as a salience parameter (see analytical arguments in Supplemental Experimental Procedures). The test would be to isolate and drive the neurons potentially corresponding to Ĉ in the brain. Activation of the Ĉ population alone should drive licking when S- recruits more activity, and should not drive licking when S- recruits less activity (**Fig. 5)**.

### Biological learning associates neural populations to different responses based on activity recruitment

Testing this prediction was impractical with our sound-based Go/NoGo discrimination protocol, because the neurons encoding common information between S+ and S- trials (Ĉ population) likely code for multiple cues, including (i) the overlap of S+ and S-representations and (ii) all cues related to the decision to visit the lick port, and cannot be isolated. Therefore, we decided to test the model predictions in an artificial but better controlled experiment in which head-fixed mice had to discriminate optogenetically driven cortical ensembles. To do so we used a custom-made setup, based on a video projector^40^, to project precise 2D light patterns through a cranial window placed above the auditory cortex in Emx1-Cre x Ai27 mice (**Fig. 6a and b**). Using intrinsic imaging, we identified the main tonotopic fields of mouse auditory cortex (**Fig. 6c**)^30,41^ and thereby reliably positioned optogenetic stimulation spots in homologous regions across mice (**Fig. 6c**). We defined three circular optogenetic stimulation spots out of which we constructed two stimuli. One of the three spots, the Ĉ spot, was common to the two S+ and S- stimuli and the two other spots corresponded to the stimulus-specific neuronal populations Ŝ+ and Ŝ- (e.g. **Fig. 6b**). Thus, the cue S+ used for Go-trials consisted of simultaneous activation of Ŝ+ and Ĉ and the cue S-consisted of simultaneous activation of Ŝ- and Ĉ. Furthermore, we were able to exclude cues common to S+ and S-, related to the visit of the port because, in the head-fixed task design, mice did not initiate the trials, which occurred at random time intervals. Thus we ensured that the only cue common to the S+ and S- stimuli was the Ĉ spot. We doubled the diameter of either the Ŝ+ or the Ŝ- spot to create a difference in cortical recruitment between the two input representations as in the model. Mice were then initially trained to obtain a water reward by licking after the coincident activation of the Ŝ+ and Ĉ spot. When 80% performance was reached in this detection task, the discrimination training started. During this stage, activation of the Ĉ population continued to occur in all trials: in half of the trials together with Ŝ+ and in the other half together with Ŝ-. Mice kept licking in the presence of the S+ spot and learned, within hundreds of trials to avoid licking in the presence of the S- spot (**Fig. 6d**), reaching a steady state performance of 94.7% ± 4.5% correct trials (hit rate 93.9% ± 3.3%, false alarm rate 4.5% ± 1.3%, n=8 mice, see **Fig. 6e&f**). Importantly, in this task setting, the stringent definition of the common Ĉ population, activated during the initial motivation training, likely resulted in the systematic establishment of strong initial connections for this population at the beginning of the discrimination training, leading to homogenous durations of the learning phase (see **Fig. 6d**) independent of salience (**Figs. 4–5**).

**Figure 6:**
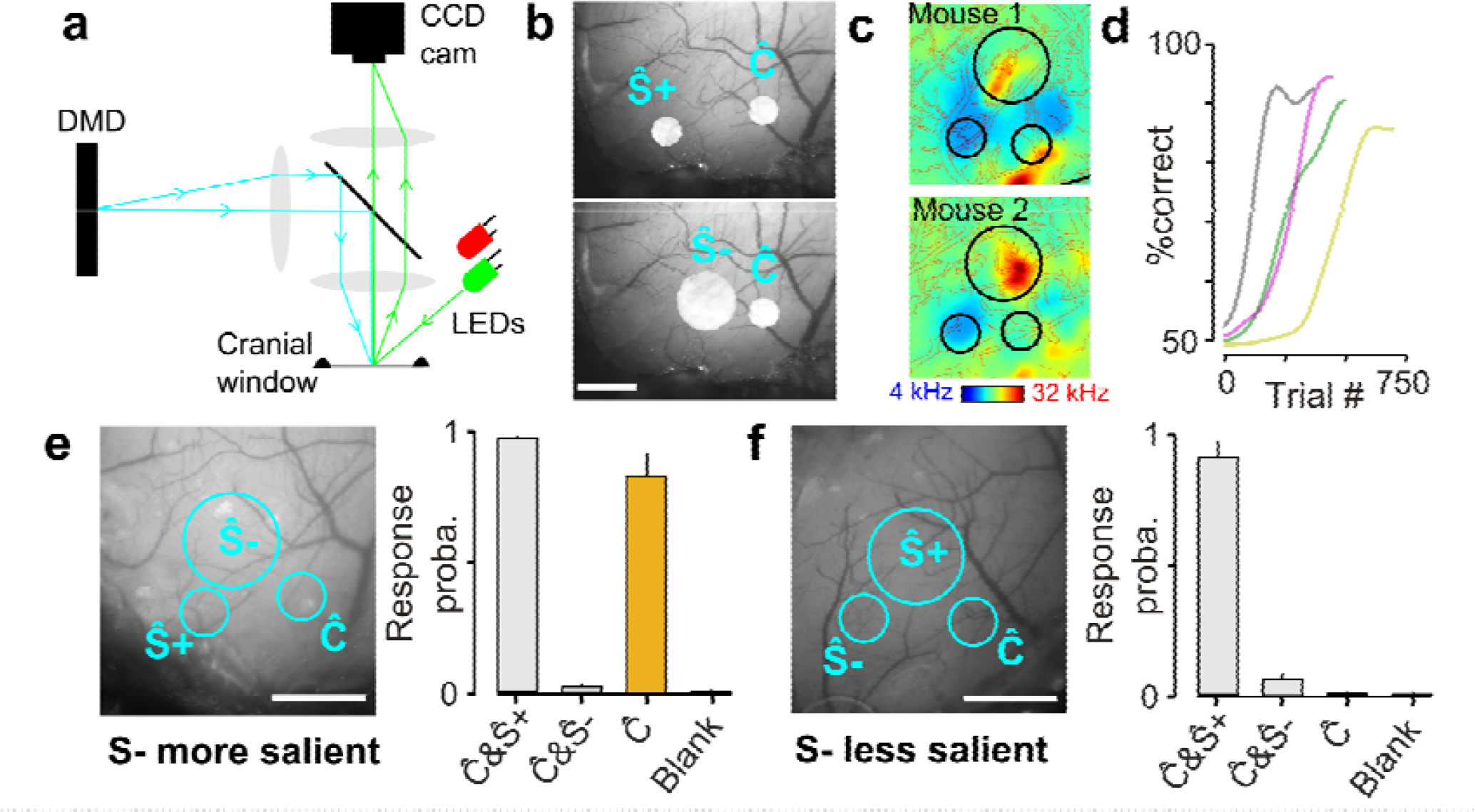
Discrimination training of multi-spot optogenetic patterns reveals a choice of learning strategy depending on the level of cortical recruitment. ***a**. Schematics of the optical setup to project arbitrary 2D light patterns onto the surface of auditory cortex. Blue light patterns from a Digital Micromirror Device (DMD) are collimated and deflected through the objective lens by a beam splitter. The surface of the cranial window can be simultaneously imaged by a CCD camera using external LEDs (green for blood vessel, red for intrinsic imaging). **b**. Examples of light patterns used for the discrimination task. Scale bars: 600 μm. **c**. Tonotopic maps obtained with intrinsic imaging for two mice, and localization of the three optogenetic stimulation spots in the same high, low and mid frequency fields. **d**. Four example learning curves for the optogenetic discrimination task. Grey: smaller Ŝ -; color: larger Ŝ -. **e**. (left) Light pattern in the task in which the S- stimulus has higher level of cortical recruitment (Ŝ- >Ŝ+). Ĉ = common component of S+ and S-stimuli. (right) Response probability on the two learnt target stimuli (Ĉ & Ŝ + vs Ĉ & Ŝ -) and on presentation of the common part of the stimulus alone in 15 catch trials (yellow). Mean ± SEM, n = 4 mice. **f**. Same as **e**, but with Ŝ + larger than Ŝ -. Mean ± SEM, n=4 mice. Scale bars: 600 μm*.

However, once mice had learned the behavioral task, we measured their response to Ĉ activation alone in catch trials that were not rewarded (catch trial probability = 0.1; 15 catch trials per mouse). In the group of mice that had a larger Ŝ- spot, we observed that activation of Ĉ elicited strong licking responses (83% ± 10% response probability, n = 4 mice, **Fig. 6e**). In contrast, in the group of mice that had a larger Ŝ+ spot, activation of Ĉ elicited no licking response (0% ± 0% licks per trials, n = 4 mice, **Fig. 6f**). These results confirm the model prediction, and demonstrate, in a causal manner, that cortical recruitment affect the choice of which stimulus is associated to a particular response. In this case, we show that the same stimulus (Ĉ activation) bearing neutral behavioral meaning is assigned by learning to be subthreshold when a larger population drives a suprathrehold response, and is assigned to be suprathreshold when a larger population drives a subthreshold response. Even if simple in essence, this result shows that cortical recruitment is a parameter influencing learning, in a manner compatible with the role of a salience parameter in reinforcement models.

## Discussion

Combining behavioral measurements, large scale two-photon imaging, optogenetics and theoretical modeling, we have shown that sounds of different quality but equal mean pressure levels can recruit highly variable levels of neuronal activity in auditory cortex, measured as the mean amount of activity in a representative subsample of neurons. We showed that cortical recruitment levels correlate with learning speed effects in a Go/NoGo task as expected if neuronal recruitment corresponds to stimulus salience. Moreover, the details of these can be precisely reproduced by a reinforcement learning model of the task. Finally, training mice to discriminate optogenetically evoked cortical patterns, and manipulating these patterns, we showed that neuronal recruitment determines which elements of the cortical representation are selected to drive each conditioned action. This corroborates, in a causal manner, the idea that cortical recruitment is a neuronal correlate of stimulus salience.

What is the origin of the differences in cortical recruitment between sounds that have equal intensity levels? In a recent study, we have suggested that nonlinearities of the auditory system select or amplify particular acoustic features such as the on- and offset features of up- and down-intensity ramps (**Fig. 2g**)^26^. This process starts as early as the cochlea which is more sensitive to some frequencies than others^42,43^ and further nonlinearities along the multiple stages of the auditory system also contribute^44^. Thus, the relative level of cortical recruitment observed for different sounds represents interesting information about central processes for acoustic feature extraction and selection. This suggests that cortical recruitment should at least partially reflect neuronal recruitment in higher subcortical areas such as thalamus. Therefore, it should be a relevant measure of salience independent of whether the discrimination task requires auditory cortex^45–47^ or does not require it^48–50^.

The idea that the amount of neuronal activity recruited by a stimulus influences behavior has been proposed in different contexts. For example, several studies indicate that attention can boost the activity of the neurons representing behaviorally relevant stimuli^6,51,52^ and thereby make it more discriminable from other stimuli^53^. Earlier reports, using direct microstimulation of the cortex, showed that low levels of neuronal recruitment can impact detection probability^54,55^. Here, we show that neuronal recruitment for stimuli that are well beyond detection threshold still impact the learning process by which they are associated to particular responses. Even if such effects are predicted by the Rescorla and Wagner model^3^, capturing their details requires a refinement of the original model. In particular, we had to introduce a more realistic learning rule in which learning speed depends not only on salience alone, but on the product of salience (modeled as neuronal recruitment) and current synaptic strength (multiplicative rule). This property has important consequences. First, it introduces variability in the relationship between salience and learning speed, through large inter-individual variations of the synaptic weights present at the beginning of the task. Second, the fact that learning speed is proportional to the product of neuronal recruitment and connectivity, allows the system to compensate weak neuronal recruitment by stronger connections (see Supplemental Experimental Procedures). This phenomenon tends to stabilize learning speed, explaining why salience does not always impact the learning phase duration, except for particular initial conditions for which compensation occurs too slowly (**Figs. 3–5**).

These complex dynamical phenomena make learning speed measurements a more complicated proxy for stimulus salience than the overshadowing protocol which relies on steady state behavior, after the dynamical phase of the association. However, it allows the comparison of salience for stimuli from the same sensory modality. Because our extended multiplicative model only diverges from the Rescorla-Wagner model for the transient dynamical part of the association process (delay and learning phases), it also reproduces overshadowing effects^24^, and more generally predicts which part of a sensory input representation is assigned to which conditioned response in a more complex task. Here, by conditioning mice to compound stimuli composed of multiple optogenetically activated neuronal ensembles of different sizes in auditory cortex (**Fig. 6**), we show, in line with reinforcement learning models, that the brain can actually establish a stimulus discrimination strategy based on the amount of activity recruited by the different subpopulations representing the stimuli.

## Acknowledgements

We thank K. Kuchibhotla, E. Harrell and M. Stüttgen for comments on the manuscript, P. Pindi, S. Sikirić and L. François for help with behavioral and imaging experiments. We thank the GENIE Project, Janelia Farm Research Campus, Howard Hughes Medical Institute, for GCAMP6s constructs. This work was supported by the Agence Nationale pour la Recherche (ANR “SENSEMAKER”), the Fyssen foundation, the DIM “Region Ile de France”, the Marie Curie Program (CIG 334581), the International Human Frontier Science Program Organization (CDA-0064-2015), by the Fondation pour l’Audition (Laboratory grant), the École Doctorale Frontières du Vivant (FdV) – Programme Bettencourt (support to AK), the DIM Cerveau et Pensée and Ecole des Neurosciences de Paris Ile-de-France (ENP, support to SC) and the Deutsche Forschungsgemeinschaft (DFG CRC1080/2).

## Author contributions

BB and SR designed the study. AK, SC, and BB performed and analyzed the imaging experiments. BB and JB performed the modeling. AD and BB performed and analyzed behavioral experiments. ZP designed the patterned optogenetic setup and SC performed the optogenetic experiments. TD designed software for data analysis and behavior. BB and SR wrote the manuscript with comments from all authors.

## Methods

### Animals

All mice used for imaging and behavior were 8 to 16 weeks old C57Bl6J and GAD2-Cre (Jax #010802) x RCL-TdT (Jax #007909) mice. Mice used for optogenetics were 8 to 16 weeks old males and female obtained by crossing homozygous Emx1^IRES-cre^ (Jax #005628) mice with Ai27 (Jax # 012567) mice to obtain expression of Td-Tomato-tagged channelrhodopsin (ChR2) in excitatory neurons of the cortex. All animal were group housed. All procedures were approved by the Austrian laboratory animal law guidelines (Approval #: M58 / 02182 / 2007 /11; M58 / 02063 / 2008 /8) and the French Ethical Committee (authorization 00275.01).

### Two-photon calcium imaging in awake mice

At least three weeks before imaging, mice were anaesthetized under ketamine medetomidine. The right *masseter* was removed and a large craniotomy (~5 mm diameter) was performed above the auditory cortex. We then performed three to five injections of 200nL (35-40nL/min), rAAV1.Syn.GCaMP6s.WPRE virus obtained from Vector Core (Philadelphia, PA) and diluted 10 times. The craniotomy was sealed with a glass window and a metal post was implanted using cyanolite glue and dental cement. At least three days before imaging, mice were trained to stand still, head-fixed under the microscope for 20 to 60 min per day. Then mice were imaged one to two hours per day. Imaging was performed using a two-photon microscope (Femtonics, Budapest, Hungary) equipped with an 8kHz resonant scanner combined with a pulsed laser (MaiTai-DS, SpectraPhysics, Santa Clara, CA) tuned at 920 nm or 900nm depending on the experiments. Images were acquired at 31.5Hz. All sounds were delivered at 192 kHz with a NI-PCI-6221 card (National Instrument) driven by Elphy (G. Sadoc, UNIC, France) through an amplifier and high frequency loudspeakers (SA1 and MF1-S, Tucker-Davis Technologies, Alachua, FL). Sounds were calibrated in intensity at the location of the mouse ear using a probe microphone (Bruel&Kjaer). In a first experiment, we played three 70 ms complex sounds at 73dB SPL preceded by two 50 ms 4kHz pure tones (inter-tone interval: 375 ms) sounds as in the behavioral task. The three sounds were played in a random order and repeated 30 times. In a second experiment, we played white noise sounds ramping -up or -down in intensity between 60dB and 85dB SPL during 2s. The ramps were repeated 20 times.

### Data analysis

Data analysis was performed using Matlab and Python scripts. Motion artifacts were first corrected frame by frame, using a rigid body registration algorithm. Regions Of Interest were selected using a semi-automated hierarchical clustering algorithm based on pixel covariance over time as described in^28^(see detailed method below). Neuropil contamination was subtracted (Kerlin et al., 2010) by apply the following equation: F_true_ (t) = F_measured_ (t) – 0.7 F_neuropil_ (t), then the change in fluorescence (∆*F*/*F_0_*) was calculated as *(F-F_0_) / F_0_*, where *F_0_* is estimated as the minimum of the low-pass filtered fluorescence over ~40 s time windows period. To estimate the time course of firing rate, the calcium signal was temporally deconvolved using the following formula*: r(t) = f’(t) + f(t) /* τ in which *f’* is the first time derivative of *f* and τ the decay constant set to 2 seconds for GCaMP6s. For the complex sounds, the population response was computed as the mean deconvolved signal across all neurons from sound onset to 500ms after sound offset. For the ramps, because the behavioural discrimination response typically occurs within few hundreds of milliseconds after the ramp onset^26^, the mean population response was evaluated from 0 to 500ms after sound onset.

### Patterned optogenetics and intrinsic imaging

To flexibly activate different activity patterns in the mouse auditory cortex, we used a computer driven (VGA input) video projector (DLP LightCrafter, Texas Instruments) which includes a strong blue LED light source (460 nm) and from which we have removed the objective. To project a two-dimensional image onto the auditory cortex surface (**Fig. 6a, b**), the image of the micromirror chip is collimated through a 150 mm cylindrical lens (Thorlabs, diameter: 2 inches) and focused through a 50 mm objective (NIKKOR, Nikon). Imaging of the cortex at the focal plane is obtained by side illumination with a green (525 nm, blood vessels) or far red (780 nm, intrinsic imaging) LED. The light collected by the objective passes through a dichroic beamsplitter (long pass, >640nm, FF640-FDi01, Semrock) and is collected by a CCD camera (GC651MP, Smartek Vision) equipped with a 50 mm objective (Fujinon, HF50HA-1B, Fujifilm). Note that the image projected to the cortical surface corresponds to a narrow cone of light extending below the surface and potentially activating ChR2 expressing neurons throughout the cortical depth. Intrinsic imaging was performed in isoflurane anesthetized mice (1.1% delivered through SomnoSuite, Kent Scientific). To compute intrinsic signal maps we divided the red light image of the cortical surface after the onset of a stimulation (average over 2 s) with 2s long pure tones (4, 8, 16 and 32kHz) by the mean image immediately before stimulus onset

### Go/NoGo discrimination behavior

Mice were water-deprived and trained daily for 200 to 300 trials. Mice first performed 4 habituation sessions to learn to obtain a water rewards (~5 µl) by licking on a spout over a threshold after the positive stimulus S+. After habituation, the fraction of collected rewards was about 80%. The learning protocol then started in which mice also received a non-rewarded, negative sound S- for which they had to decreasing licking below threshold to avoid an 8s time-out. For the freely moving complex sound discrimination, S+ and S- sounds consisted of two 4 kHz pips (50 ms) followed by one of the three 70 ms complex click shown in **Fig. 1a**. The interval between the offset and onset of the pips and click was 375 ms. Licking was assessed 0.58 sec after the specific sound cue in a 1 sec long window by an infrared beam at the spout. For the intensity ramp discrimination, licking was assessed in a 1.5s window after sound offset. In both cases, licking was considered above threshold if the infrared beam in front of the licking tube was broken during 75% of the measurement time-window. Positive and negative sounds were played in a pseudorandom order with the constraint that exactly 4 positive and 4 negative sounds must be played every 8 trials. For learning curves, performance was measured as the fraction of correct positive and correct negative trials over bins of 100 trials. For the optogenetically-driven, head-fixed discrimination task, the S+ and S- stimuli were each composed of two disks of blue light (465 nm) flashing at 20 Hz for 1 s. One of the two disks (noted Ĉ) was common to S+ and S-stimuli, the other disk was condition-specific. The three disk locations were chosen in similar tonotopic locations across mice based on intrinsic imaging maps. They were precisely repositioned for every training session using an automated registration procedure based on blood-vessel patterns. A strong masking light was used to prevent the animal from using visual cues in the task. The common disk was 360 µm in diameter. In one set of mice, the disk specific to S- was 720 µm in diameter, while the S+ specific disk was 360 µm in diameter. In the other set of mice, sizes of the specific disks were swapped. Head-fixed mice performed 200 to 300 trials per day with an inter-trial interval randomized between 3 s and 7 s. Individual licks were detected through an electric circuit connecting the mouse and the lick tube. Then, each trial was started only if the mouse was not spontaneously licking for at least 3 s (in3 addition to the inter-trial interval). Mice were first trained to respond to the S+ stimulus by producing at least 3 to 5 licks (depending on the mouse) to get the 5 µL water reward. When the mouse could collect more than 80% of the rewards, the S- stimulus was introduced. Licking above threshold after S- was punished with a 7 s timeout.

### Reinforcement learning model

The model has been described extensively in a previous publication^24^. In short, it is composed of three sensory units (Ŝ+, Ŝ- and Ĉ, representing populations of neurons) whose activity described by a three-dimensional vector *x*⃗ and which are connected to a simple decision circuit (**Fig. 4a**). Cortical recruitment is modeled by changing the firing value of the sound units. When the S- stimulus recruits less activity than the S+ stimulus, the input vectors are: *x*⃗ _*S*+_ = [1 0 2] or *x*⃗_*S*−_ = [1 1 0]. When S- recruits more activity than S+, the input vectors are: *x*⃗ _*S*+_ = [1 0 1] or *x*⃗ _*S*−_ = [1 2 0].

The decision circuit is composed of all-or-none response unit (*y* = 0 or 1) which linearly sums the three sensory inputs (representing synaptic populations) under the form of three direct excitatory connections and of a graded feed-forward inhibition from a virtual inhibitory unit in fact equivalent to three direct inhibitory connections. The output of model is described by a single equation for the decision unit:

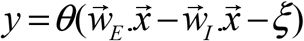

in which *θ* is the Heaviside step function. *w*⃗_*E*_ and *w*⃗_*I*_ are three-dimensional positive vectors describing the excitatory synaptic weights from the sensory units to the decision and inhibitory units respectively. The variable *ξ* is a Gaussian random noise process of unit variance which models the stochasticity of behavioral choices.

Based on the action outcome (*R* = 1 for a reward, *R* = −1 for no reward), the learning rule for the synaptic weights is implemented as:

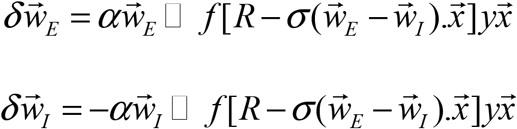

in which  is the Hadamard (element-wise) product implementing the multiplicative rule, *yx*⃗ is a Hebbian term, *α* is the learning rate and *σ* is a parameter related to the noisiness of the model and setting its asymptotic performance. To account for the faster improvement of performance for rewarded as compared to non-rewarded trials, positive expectation errors are more strongly weighted than negative ones, thanks to the asymmetric function *f[u]* = *u* if *u* ≤ 0 and *f[u] = νu* if *u* > 0. The parameter ν is typically larger than 1, consistent with the activity of basal ganglia dopaminergic neurons in mice^56^ and monkeys^57^ coding for reward expectation error.

As described above, the equations of the model are stochastic due to the Gaussian random noise process *ξ*. To compute the response probability estimates plotted throughout the study, we used a previously established probability equation (Bathellier et al., 2013), valid for learning dynamics much slower than fluctuations (ergodic approximation). The probability to make a lick response given the input vector *x*⃗ _*S*+_ or *x*⃗ _*S*−_ is:

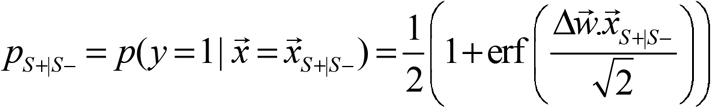

where Δ*w*⃗= *w*⃗_*E*_ – *w*⃗_*I*_ represents now the average observed values of difference between the excitatory and inhibitory connections. In addition, the plasticity equations become:

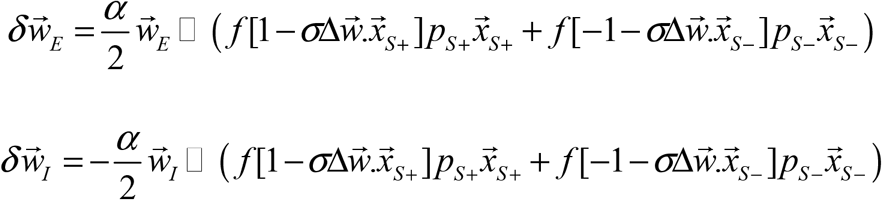

### Statistical tests

Unless otherwise specified, all quantifications are given as mean ± standard error (SEM). To statistically assess the differences between paired measurements (e.g. activity for two different sounds elicited in the same neuronal populations) we used the non-parametric Wilcoxon signed rank test. To compare two sets of measurements (e.g. delay and learning phase duration for two groups of mice) we used the non-parametric Wilcoxon rank sum test. Assessment of the differences in the fraction of responsive neurons for different sounds was done with the χ^2^ test which evaluates differences in the distributions of two binary variables.

### Data and software availability

Datasets, analysis software and codes for running the simulations of our model are available upon request to Brice Bathellier.

